# Towards the design of multiepitope-based peptide vaccine candidate against SARS-CoV-2

**DOI:** 10.1101/2020.07.07.186122

**Authors:** Hasanain Abdulhameed Odhar, Salam Waheed Ahjel, Suhad Sami Humadi

**Author notes:** Corresponding author: Hasanain Abdulhameed Odhar, Department of pharmacy, Al-Zahrawi University College, Karbala, Iraq.

## Abstract

Coronavirus disease 2019 is a current pandemic health threat especially for elderly patients with comorbidities. This respiratory disease is caused by a beta coronavirus known as severe acute respiratory syndrome coronavirus 2. The disease can progress into acute respiratory distress syndrome that can be fatal. Currently, no specific drug or vaccine are available to combat this pandemic outbreak. Social distancing and lockdown have been enforced in many places worldwide. The spike protein of coronavirus 2 is essential for viral entry into host target cells via interaction with angiotensin converting enzyme 2. This viral protein is considered a potential target for design and development of a drug or vaccine. Previously, we have reported several potential epitopes on coronavirus 2 spike protein with high antigenicity, low allergenicity and good stability against specified proteases. In the current study, we have constructed and evaluated a peptide vaccine from these potential epitopes by using in silico approach. This construct is predicted to have a protective immunogenicity, low allergenicity and good stability with minor structural flaws in model build. The population coverage of the used T-cells epitopes is believed to be high according to the employed restricted alleles. The vaccine construct can elicit efficient and long-lasting immune response as appeared through simulation analysis. This multiepitope-based peptide vaccine may represent a potential candidate against coronavirus 2. However, further in vitro and in vivo verification are required.

## Background

In December 2019, multiple pneumonia cases of unknown etiology were reported in Wuhan, China **[1]**. Later on, genomic analysis of samples collected from admitted patients had revealed that a novel beta-coronavirus was the causative pathogen **[2]**. This RNA virus was temporarily known as 2019 novel coronavirus (2019-nCoV), but was renamed later as severe acute respiratory syndrome coronavirus 2 (SARS-CoV-2) **[3]**. The disease caused by this virus is known as coronavirus disease 2019 (COVID-19), it is usually characterized by fever, dry cough, fatigue and muscles pain. Additionally, dyspnea can be observed in some COVID-19 patients and it may progress into acute respiratory distress syndrome (ARDS) **[4]**. The transmission of SARS-CoV-2 is largely dependent on respiratory droplets generated through sneezing or coughing **[5]**. The median incubation period of SARS-CoV-2 is estimated to be 5.1 days with 95% confidence interval of 4.5 to 5.8 days **[6]**. The outbreak of SARS-CoV-2 was recognized as a global pandemic threat on March 11, 2020 **[7]**.

SARS-CoV-2 has a genomic sequence similarity of 79% with a previously known coronavirus, SARS-CoV **[8]**. Both viruses can invade host alveolar cells through interaction of viral spike protein with angiotensin converting enzyme 2 (ACE2). However, the binding affinity of SARS-CoV-2 to ACE2 seems to be 10-20 times higher **[9]**. It is believed that COVID-19 pathogenesis may involve blockade of ACE2 and subsequent imbalance between angiotensin- (1-7) and angiotensin II. This suggests that ACE2 may be further involved in COVID-19 pathogenesis beyond viral entry point **[10]**.

Similar to SARS-CoV, no specific treatment is currently available to combat COVID-19. However, management of ARDS does involve the use of oxygen-based therapy and antibiotics against possible sepsis **[11]**. In attempts to fight COVID-19, clinical trials are keep going to assess the effect of several antiviral agents like Remdesivir **[12]**. Additionally, computational modelling attempts have proposed the repurposing of several FDA approved drugs against SARS-CoV-2 **[13,14]**.

In the same time, no vaccine was ever developed for any coronavirus. However, advanced techniques had been employed to generate potential SARS-CoV-2 vaccine candidates like mRNA based vaccine, viral vector vaccine and viral subunit vaccine **[15]**. Recently, immuno-informatics tools had been used to screen SARS-CoV-2 proteins for potential B-cells and T-cells epitopes **[16,17]**. The prediction of these epitopes had enabled virtual design of peptide based vaccine against SARS-CoV-2 **[18]**.

Previously, we have predicted several potential epitopes within SARS-CoV-2 spike protein sequence. Of interest was the linear B-cells epitope with sequence ‘GFNCYFPLQSYGF’, this epitope is believed to be part of spike protein receptor binding protein (RBD) that is involved in interaction with ACE2 **[17,19]**. In the current study, we have constructed a peptide-based vaccine design by combining seven potential epitopes predicted from our previously published findings **[17]**. Then, we have evaluated this vaccine design by using several immuno-informatics tools to affirm its stability, safety and efficiency. The aim of this study is to present a multiepitope-based peptide vaccine design for potential use against SARS-CoV-2.

## Methodology

### Setting up a research plane

A flowchart that summarizes the steps of vaccine construction and evaluation study can be seen in **Figure 1**.

**Figure 1:**
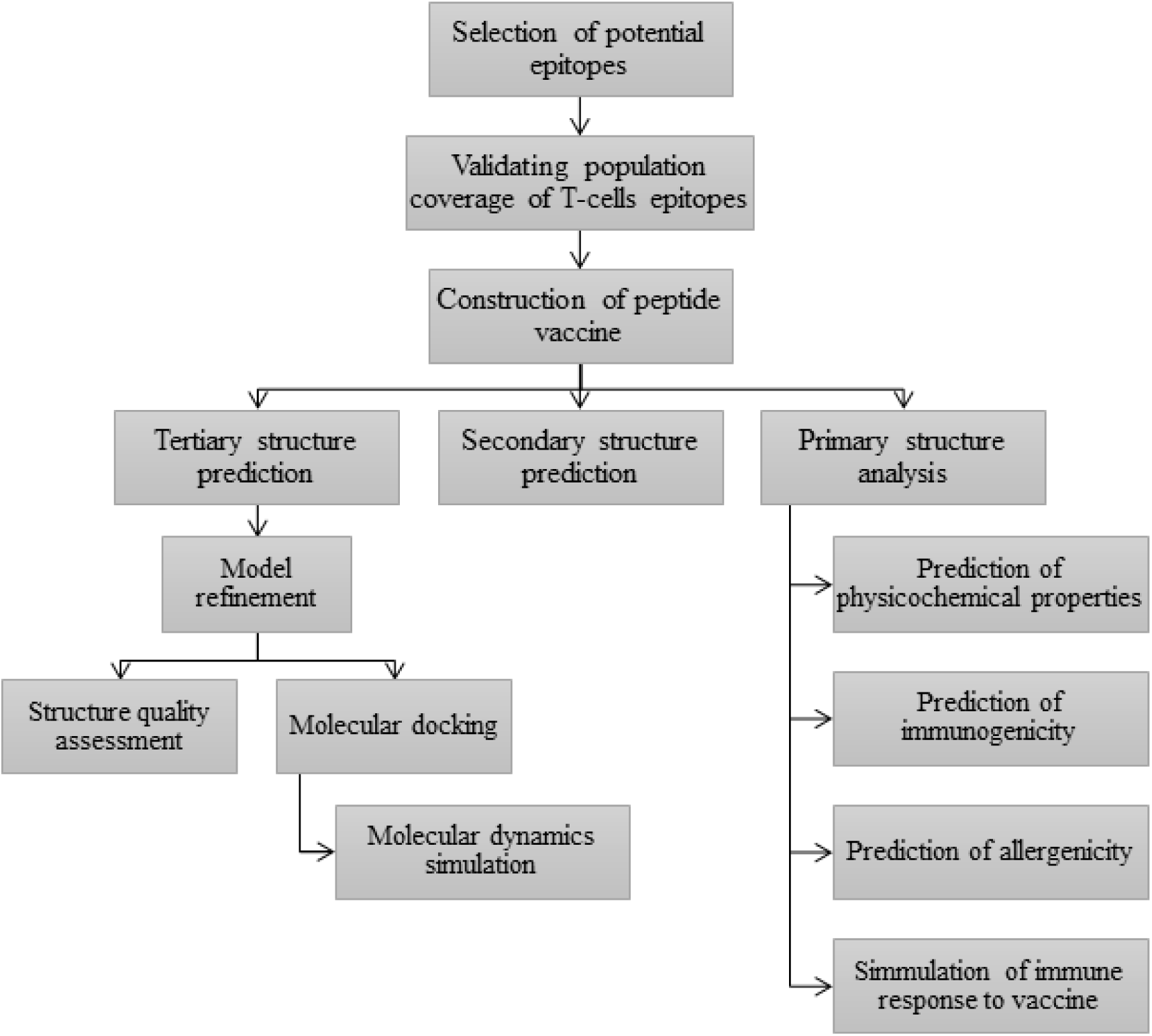
A flowchart for vaccine construction and assessment steps.

### Selection of potential epitopes for peptide vaccine build

We have selected seven potential epitopes for the construction of vaccine candidate. The sequence of these epitopes can be seen in **Table 1**, they have high antigenicity, low allergenicity and good stability against selected proteases as predicted by our previously published study **[17]**.

**Table 1:**
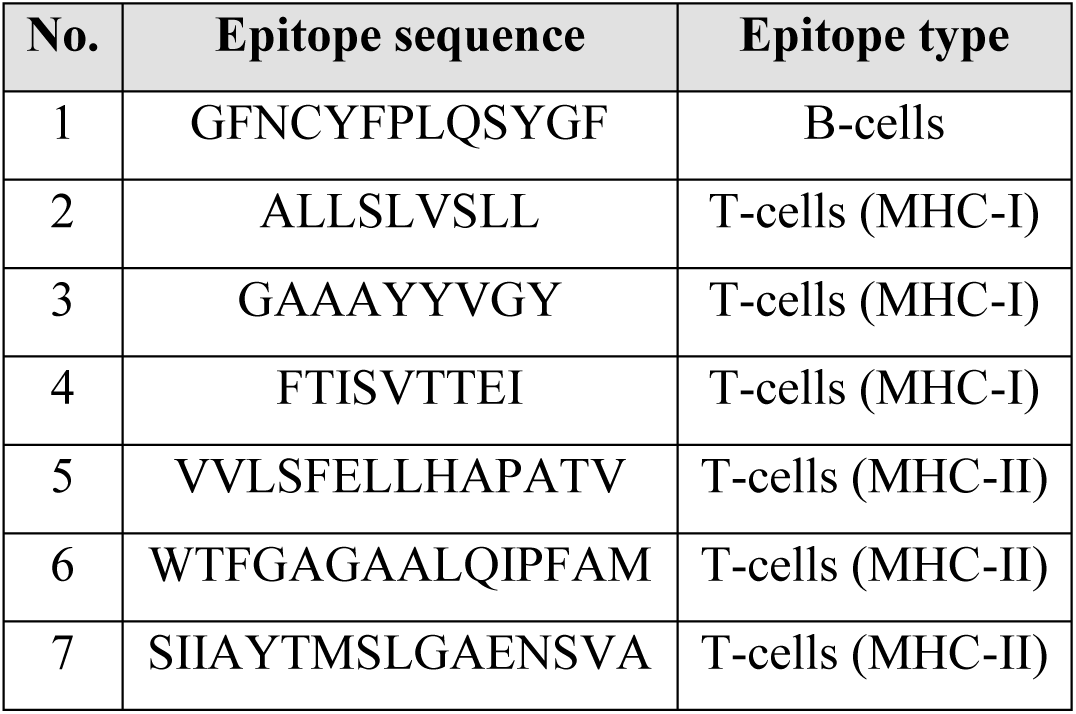
Linear epitopes previously predicted on SARS-CoV-2 spike protein crystal.

### Population coverage of selected T-cells epitopes

Major histocompatibility complex (MHC) molecule are highly polymorphic and expressed at different frequencies in different populations. A successful peptide-based vaccine should have multiple epitopes with different human leucocyte antigen (HLA) binding specificities in order to increase population coverage. We have used population coverage tool, available through the immune epitope database (IEDB), to predict the response of population in different locations of the world to the selected six T-cells epitopes. This web-based tool can predict the response to specific epitope by using MHC binding and/or T-cells restriction data and also HLA genotypic frequencies **[20,21]**. We have specified the number of T-cells epitopes to 6 peptides, then different areas around the world were selected for population coverage screening. Class I and II combined option was used to account for both MHC class I and MHC class II restricted T-cells epitopes. For MHC class I epitopes, we have used the following restricted alleles: A*01:01, A*02:01, A*02:06, A*03:01, B*07:02, A*11:01, A*23:01, A*26:01, B*35:01 and B*51:01. While for MHC class II epitopes, we have selected the following 10 alleles: DRB1*01:01, DRB1*03:01, DRB1*04:01, DRB1*04:04, DRB1*04:05, DRB1*07:01, DRB1*09:01, DRB1*11:01, DQB1*02:01 and DQB1*03:01.

### Peptide vaccine construction

After population coverage analysis, these seven epitopes were then merged together in a sequential manner by using AAY linker. This linker is considered as a cleavage site of mammalians proteasomes and can increase epitopes presentation by enhancing the formation of natural epitopes and reducing the formation of junctional epitopes **[22]**. To enhance immunogenicity of vaccine design, we have linked a human beta-defensin 2 chain A as an adjuvant. This adjuvant is considered an antimicrobial peptide with a length of 41 amino acids, it was linked to the N-terminus of vaccine design by using EAAAK linker **[23,24]**. The EAAAK linker has a rigid nature and can connect the adjuvant to the peptide vaccine with adequate separation and minimal intervention **[25]**.

### Prediction of physicochemical properties, immunogenicity and allergenicity of vaccine design

We have used ProtParam web-based tool to predict different physical and chemical characteristics of vaccine construct **[26]**. To assess immunogenicity, we have employed VaxiJen v2.0 predictive tool with a threshold value greater than 0.5. VaxiJen v2.0 tool can predict protective antigens with no need for sequence alignment, it depends mainly on physicochemical properties of the submitted peptide **[27]**. Then we have used AllergenFP v.1.0 tool to evaluate allergenicity potential of vaccine sequence **[28]**. The sequence of peptide vaccine was submitted for these tools as one letter format.

### Secondary and tertiary structures prediction of vaccine design

The secondary structure of vaccine construct was predicted by using PSIPRED 4.0 tool **[29]**. While tertiary structure was modelled by using SPARKS-X, this web-based tool can recognize protein folding through sequence alignment **[30]**. For these prediction tools, the sequence of vaccine design was submitted as one letter code. To further improve vaccine tertiary structure, the generated PDB file was then submitted to Galaxy refine server. This refinement server applies molecular dynamics (MD) simulation to implement repeated cycles of structural perturbation with subsequent relaxation **[31]**.

### Vaccine structure validation

The peptide vaccine structure was submitted as PDB file to RAMPAGE server for Ramachandran plot analysis. This analysis provides a view for protein conformation by plotting the torsional angles for each residue **[32]**. Then, the vaccine PDB was assessed by ProSA-web tool. This validation software can generate an energy plot and calculate a quality score for each submitted protein. If the calculated score is outside a range specific for native proteins, then the submitted peptide may have structural flaws **[33]**. Finally, the distribution pattern of the different atoms in the peptide vaccine was evaluated by using ERRAT server. This web-based tool can statistically analyze the non-bonded interactions between various atom types in the submitted model build **[34]**.

### Molecular docking and dynamics simulation study

The vaccine PDB was then docked into Toll-like receptor 8 (TLR8) crystal by using PatchDock web server. This web-based server uses shape complementarity algorithm for protein-protein docking **[35]**. We have used only chain A of TLR8 crystal with PDB code 5Z14, water molecules and other bounded chemicals were removed from the crystal by using UCSF chimera version 1.13.1 **[36,37]**. The Toll-like receptor is very important in linking adaptive immunity with innate immunity. It plays an essential role in initiating host immune response through upregulation of inflammatory cytokines. TLR8 has the ability to detect viral single-stranded RNA **[38,39]**. The generated docking complex with the highest geometric shape complementarity score was then downloaded as PDB file and visualized by using PyMOL version 2.3 and DIMPLOT **[40,41]**.

Later, this docking complex was subject to molecular dynamics (MD) analysis by using YASARA Dynamics version 19.12.14 **[42]**. During dynamics simulation, the various atoms and molecules of protein-protein complex are allowed to interact and move. The trajectories of these atoms and interaction potential energy are determined for a specific period of time **[43]**. MD analysis was carried out for 10 nanoseconds by using the same protocol applied in our previously published article **[14]**.

### Simulation of immunogenic response to vaccine construct

The ability of peptide vaccine to activate various components of host immune system was predicted by using C-ImmSim server. This modelling platform combines the dynamics of immune system together with genomic information **[44]**. At first, the vaccine sequence was submitted in FASTA format with one letter code. Each one-time step during simulation is equivalent to 8 hours of real time, the simulation steps were adjusted to 1000 in order to allow immune response modelling for about 333 days. During the simulation, three injections of the vaccine were applied to generate efficient and long-lasting immune response. Each injection contains 1000 units of vaccine construct. The time steps for these injections were specified as 1, 84 and 168 which is equivalent in real time to 1, 28 and 56 days respectively. It is recommended to apply a time frame of at least one month between vaccine priming dose and subsequent boost dose in order to allow sufficient time for memory cells proliferation and development **[45,46]**. We have used other simulation parameters with default values.

## Results and discussion

The selected six T-cells epitopes showed high predicted population coverage according to MHC restricted alleles. According to **Figure 2**, the worldwide coverage of these epitopes was 97.68%. The lowest population coverage was reported in Central America with a percentage of 60.2, while the highest coverage with 99.14% was recorded for Europe area. The population coverage data in **Figure 2** indicates that the selected T-cells epitopes are suitable for construction of peptide vaccine and can minimize MHC restriction of T-cell responses.

**Figure 2:**
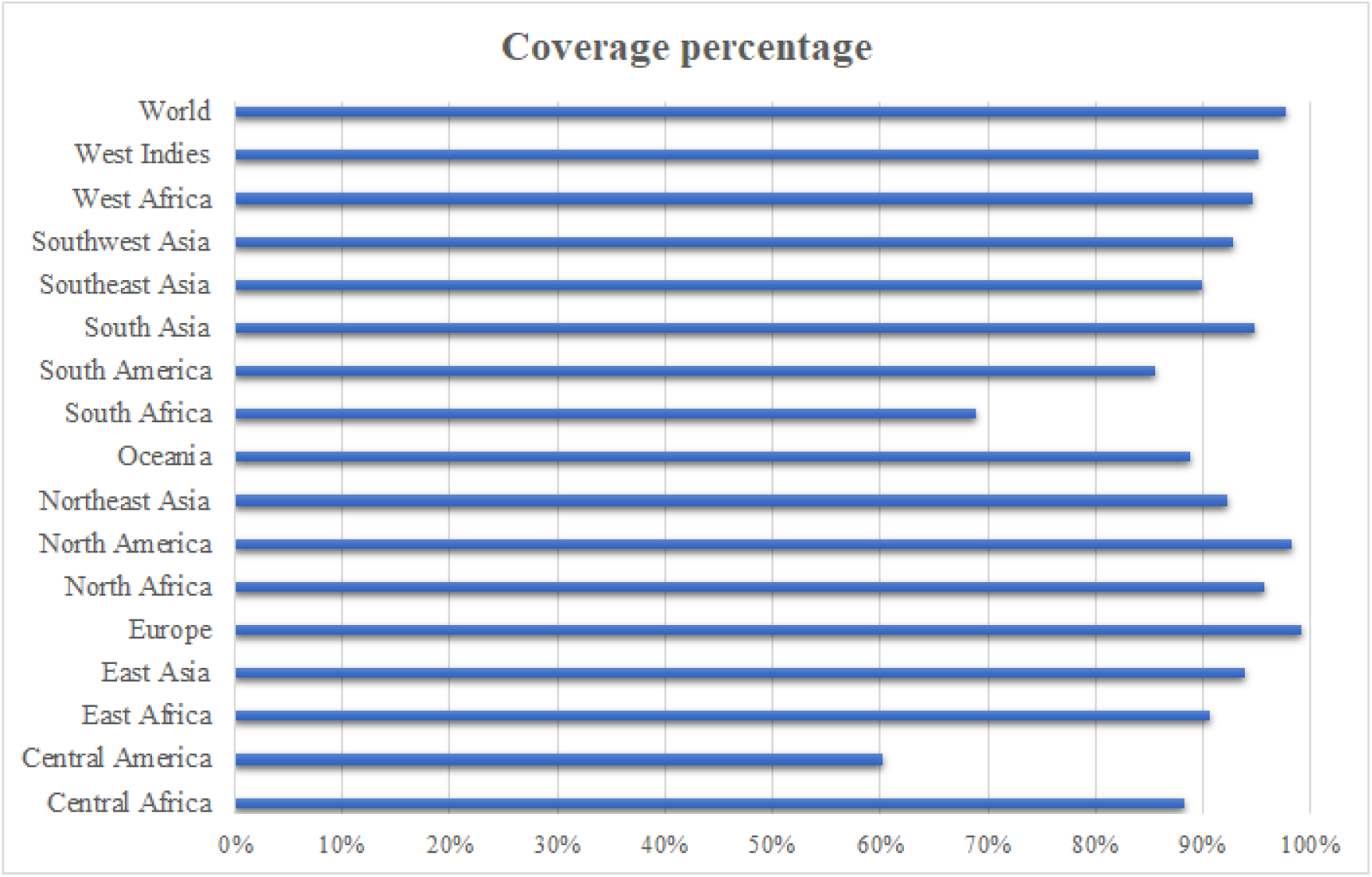
Population coverage percentage for selected T-cells epitopes according to MHC restricted alleles.

The primary sequence of peptide vaccine was constructed by merging the seven potential epitopes together by using AAY linker, while the adjuvant human beta-defensin 2 chain A was linked to the N-terminus of vaccine construct by using the rigid linker EAAAK. The primary structure of peptide vaccine can be seen in **Figure 3 (A)** as one letter format. Prediction of vaccine secondary structure as appeared in **Figure 3 (B)** revealed that the percentage of β-strand, α-helix and coil are 10, 53 and 37 respectively. The tertiary structure of vaccine design was modelled by using SPARKS-X server with a Z-score of 5.26, the model was further refined by using Galaxy refine server with a root mean square deviation (RMSD) of 0.496 from initial build. The refined tertiary structure of peptide vaccine can be seen in **Figure 3 (C)**.

**Figure 3:**
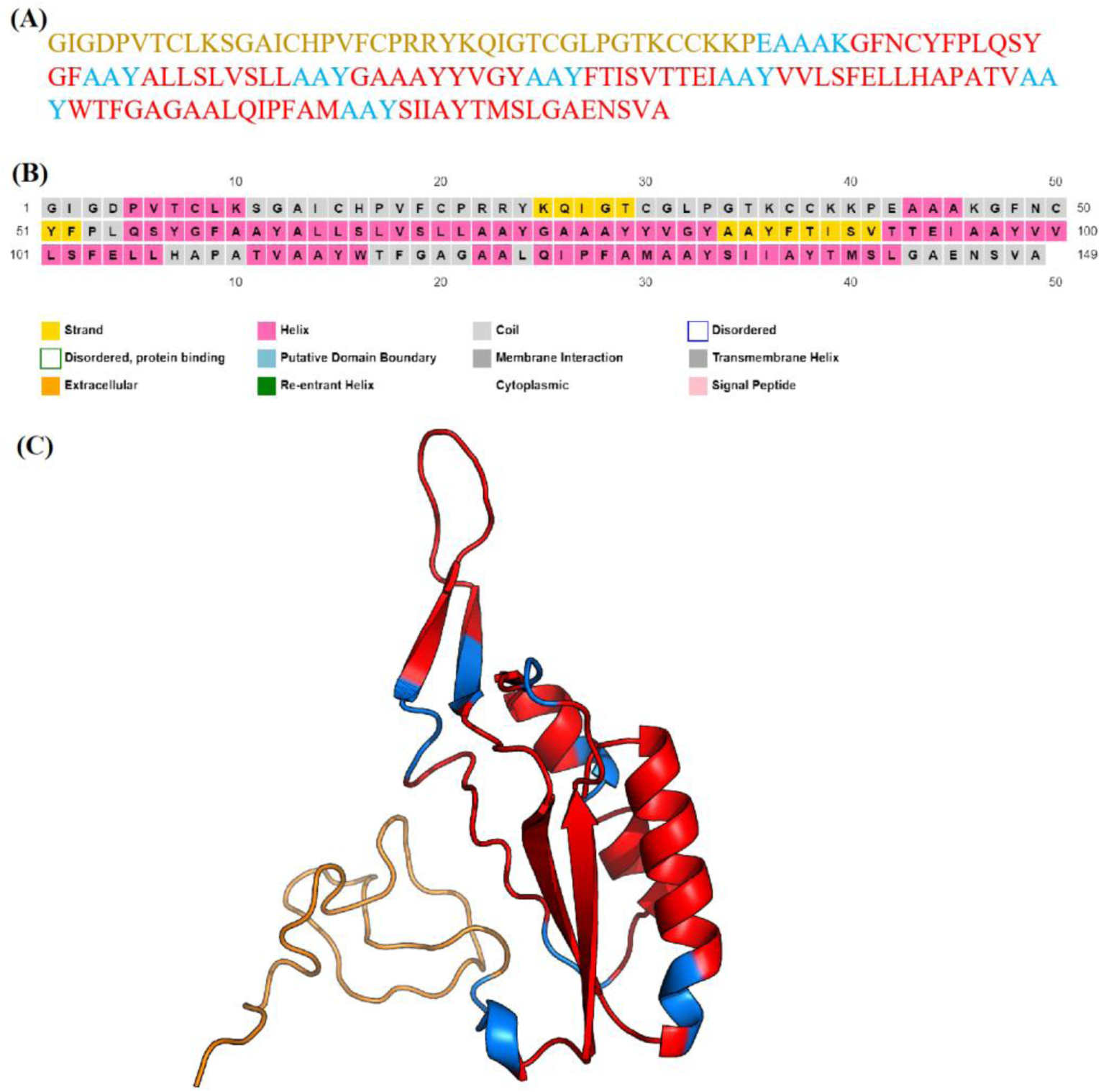
**(A)** The primary structure of vaccine construct is presented as one letter format, the sequence of adjuvant, linkers and epitopes are colored by gold, light blue and red respectively. **(B)** Prediction of vaccine design secondary structure, where β-strand, α-helix and coil are colored as yellow, pink and gray respectively. **(C)** The refined model of peptide vaccine as a tertiary structure, image was generated by using PyMOL version 2.3 **[40]**. The adjuvant, linkers and epitopes are also colored by gold, light blue and red respectively.

A summary of predicted physicochemical properties, immunogenicity and allergenicity of vaccine construct can be observed in **Table 2**. According to this table, the peptide vaccine seems to have a net positive charge as the number of positively charged residues exceeds the number of negatively charged ones. Also, the isoelectric point (IP) is reported to be 8.46. An isoelectric point value greater than 7 indicates that the overall charge of the peptide is positive. The isoelectric point is defined as the PH value of a solution at which the protein possesses zero net charge **[47]**. The calculated instability index (II) of peptide vaccine is 35.39, proteins with instability index below 40 are considered stable **[48]**. The peptide vaccine half-life is anticipated to be 30 hours in mammalian reticulocytes and greater than 10 hours in Escherichia coli. The peptide vaccine seems to be a probable antigen with antigenicity score of 0.7432 and it is probably non-allergenic.

**Table 2:**
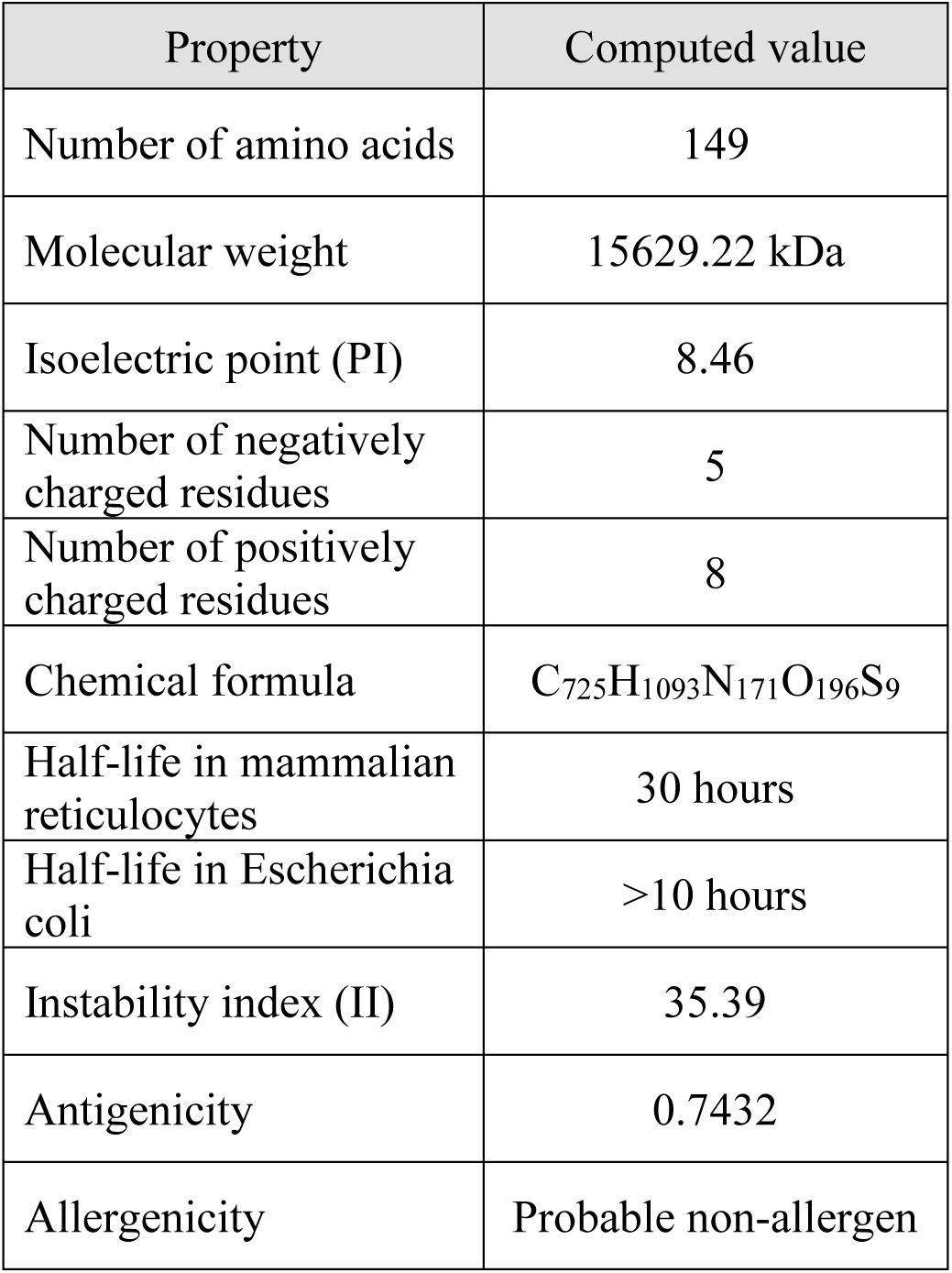
Predicted physical and chemical characteristics of peptide vaccine along with its antigenicity and allergenicity potentials.

| Property | Computed value |
| --- | --- |
| Number of amino acids | 149 |
| Molecular weight | 15629.22 kDa |
| Isoelectric point (PI) | 8.46 |
| Number of negatively charged residues | 5 |
| Number of positively charged residues | 8 |
| Chemical formula | C<sub>725</sub>H<sub>1093</sub>N<sub>171</sub>O<sub>196</sub>S<sub>9</sub> |
| Half-life in mammalian reticulocytes | 30 hours |
| Half-life in *Escherichia coli* | >10 hours |
| Instability index (II) | 35.39 |
| Antigenicity | 0.7432 |
| Allergenicity | Probable non-allergen |

The validation results of vaccine refined tertiary structure can be observed in **Figure 4**. Based on analysis of Ramachandran plot in **Figure 4 (A)**, 93.2% of residues are in favored position of torsional angles plot while 4.8% are in allowed region. Only 2.0% of the residues in the refined model are considered outliers. Then, the Z-score of the refined model was recorded to be -3.59 by using ProSA-web tool. This quality score is located within the Z-scores range calculated for native proteins of similar size as can be seen in **Figure 4 (B)**. The model quality was also assessed, as presented in **Figure 4 (C)**, by plotting predicting energy as a function of sequence position. The plot for window size of 40 residues is smoother than 10 residues size. As obvious, our refined model may have some structural flaws as the predicted energy for window size of 40 residues jumps slightly higher than zero in part of the plot. Finally, the overall quality score of ERRAT server is reported to be 82.645. This score represents the percentage of protein sequence with predicted error value that falls below rejection limit. A plot of error value versus residues number can be seen in **Figure 4 (D)**. All the above structural validation results predict that the refined construct has a good stability with minor flaws.

**Figure 4:**
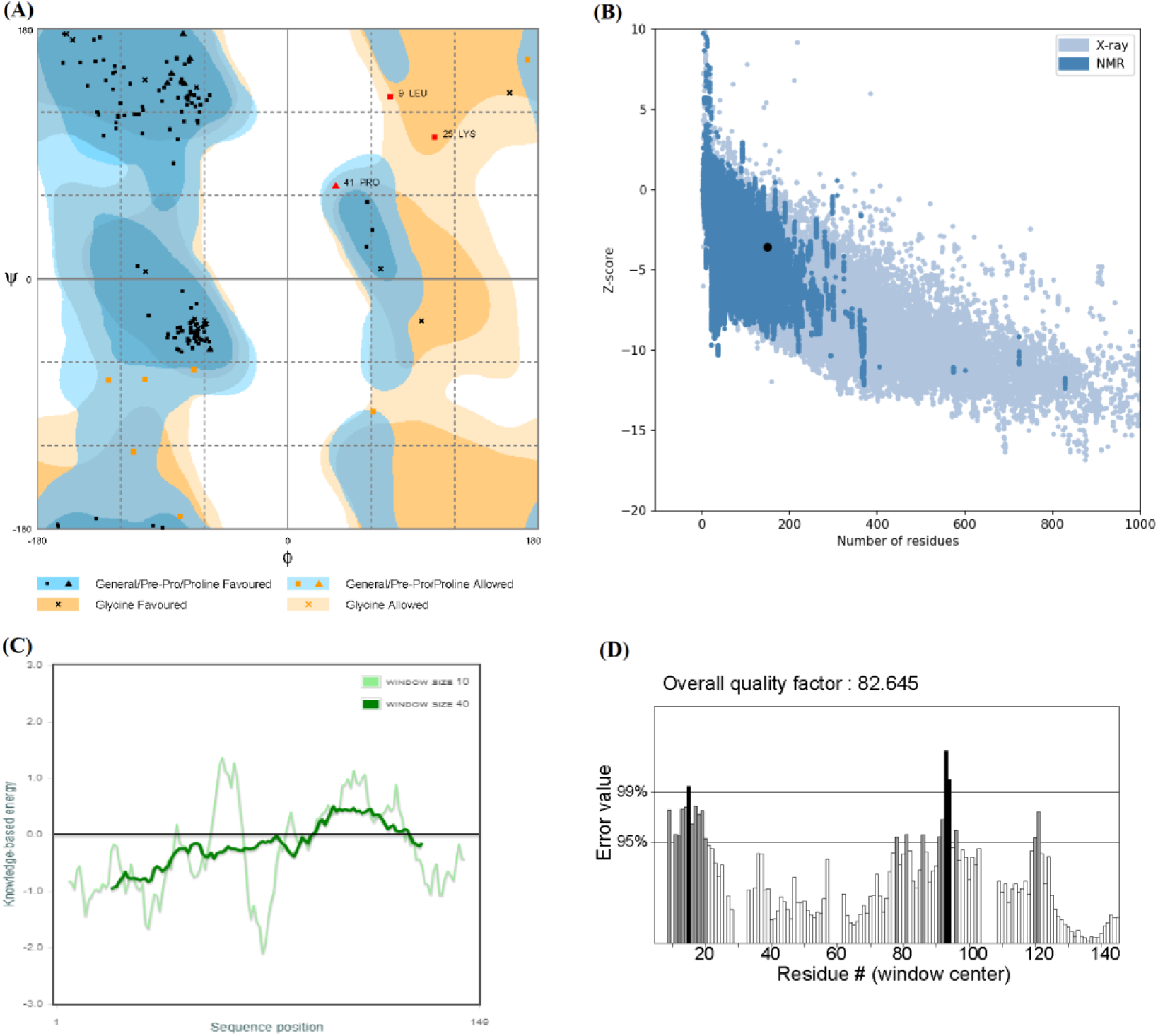
**(A)** Ramachandran plot analysis for refined vaccine construct. **(B)** Quality assessment by plotting Z-score as a function of protein size. **(C)** Model evaluation by plotting energy versus residues position. **(D)** A plot for predicted error value for each residue in vaccine candidate, the 99% and 95% lines represent confidence interval for rejection limit.

Molecular docking analysis of vaccine construct against TLR8 had generated many potential complexes. An overview for docking result of the first ranked complex can be seen in **Figure 5**. This vaccine-TLR8 complex has a geometric shape complementarity score of 17544 with approximate interface area of 2745.70. A cartoon representation for this complex is well presented in **Figure 5 (A)**. According to analysis of protein-protein interface illustrated in **Figure 5 (B)**, The peptide vaccine was able to form 5 hydrogen bonds with TLR8 interface residues. The anticipated length of these bonds is less than 3.5 Angstrom.

**Figure 5:**
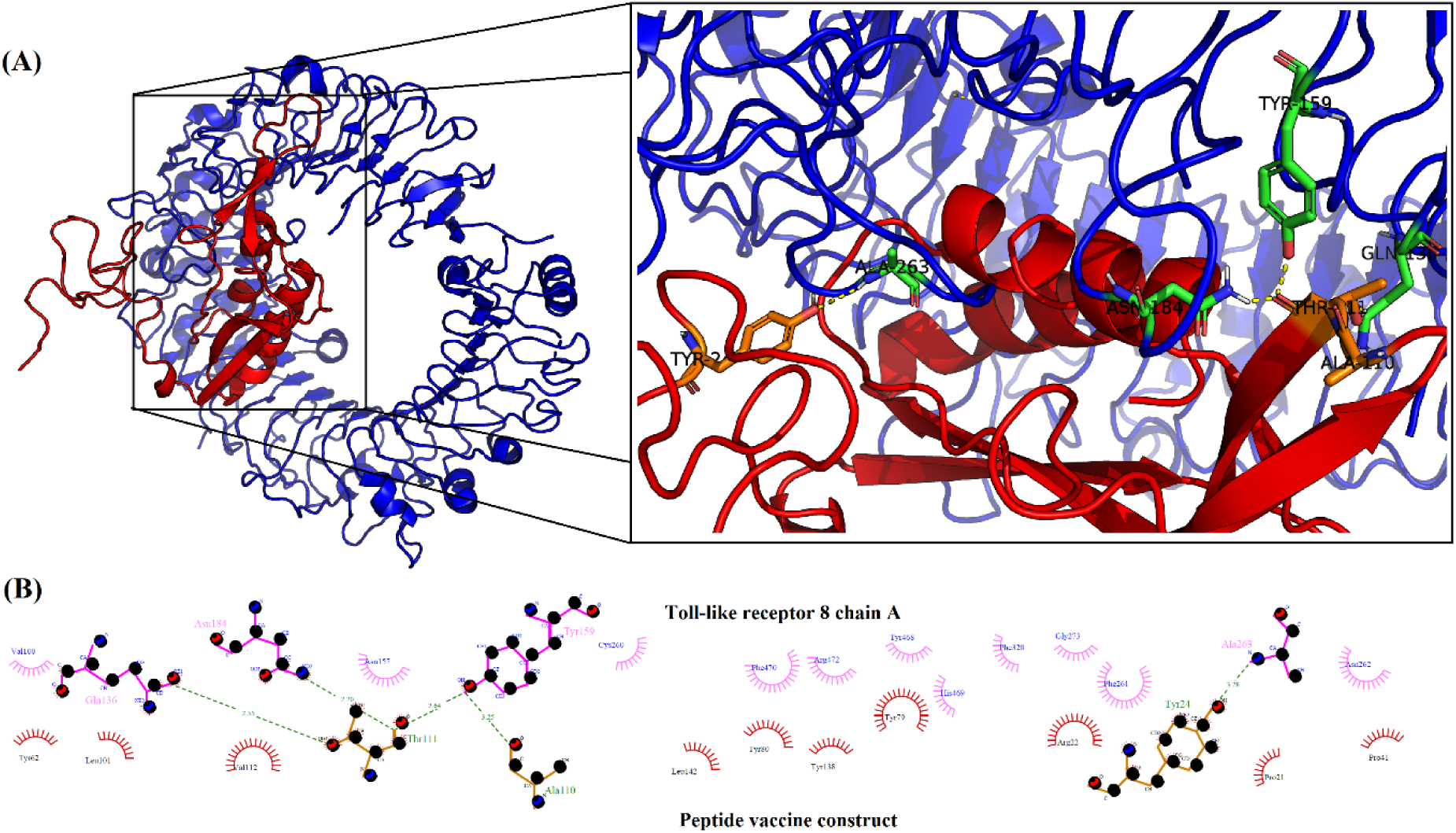
**(A)** A cartoon illustration for first ranked docking complex of peptide vaccine and TLR8. The peptide vaccine is colored by red and its interacting residues at the interface are colored by orange. On the other hand, TLR8 is colored by blue and its residues involved in interface interaction are colored by green. The hydrogen bonds are represented by yellow dashed lines. **(B)** A two-dimensional representation for vaccine-TLR8 interface. Hydrogen bonds are illustrated as green dashed lines while the small multiple lines refer to hydrophobic interactions between interface residues.

Molecular Dynamics (MD) analysis results for the first ranked docking complex of vaccine and TLR8 can be seen in **Figure 6**. Based on total potential energy calculations in **Figure 6 (A)**, the complex looks to be stable with minimal changes in potential energy predicted throughout simulation period. However, the vaccine construct seems to exhibit a higher flexibility per residue than TLR8 as noted from root mean square fluctuation (RMSF) calculations in **Figure 6 (B)**. The residues of peptide vaccine show a higher variation in position from its simulation time averaged position (reference position). The results of both molecular docking and dynamics simulation may indicate the preferential binding of vaccine design to Toll-like receptor 8.

**Figure 6:**
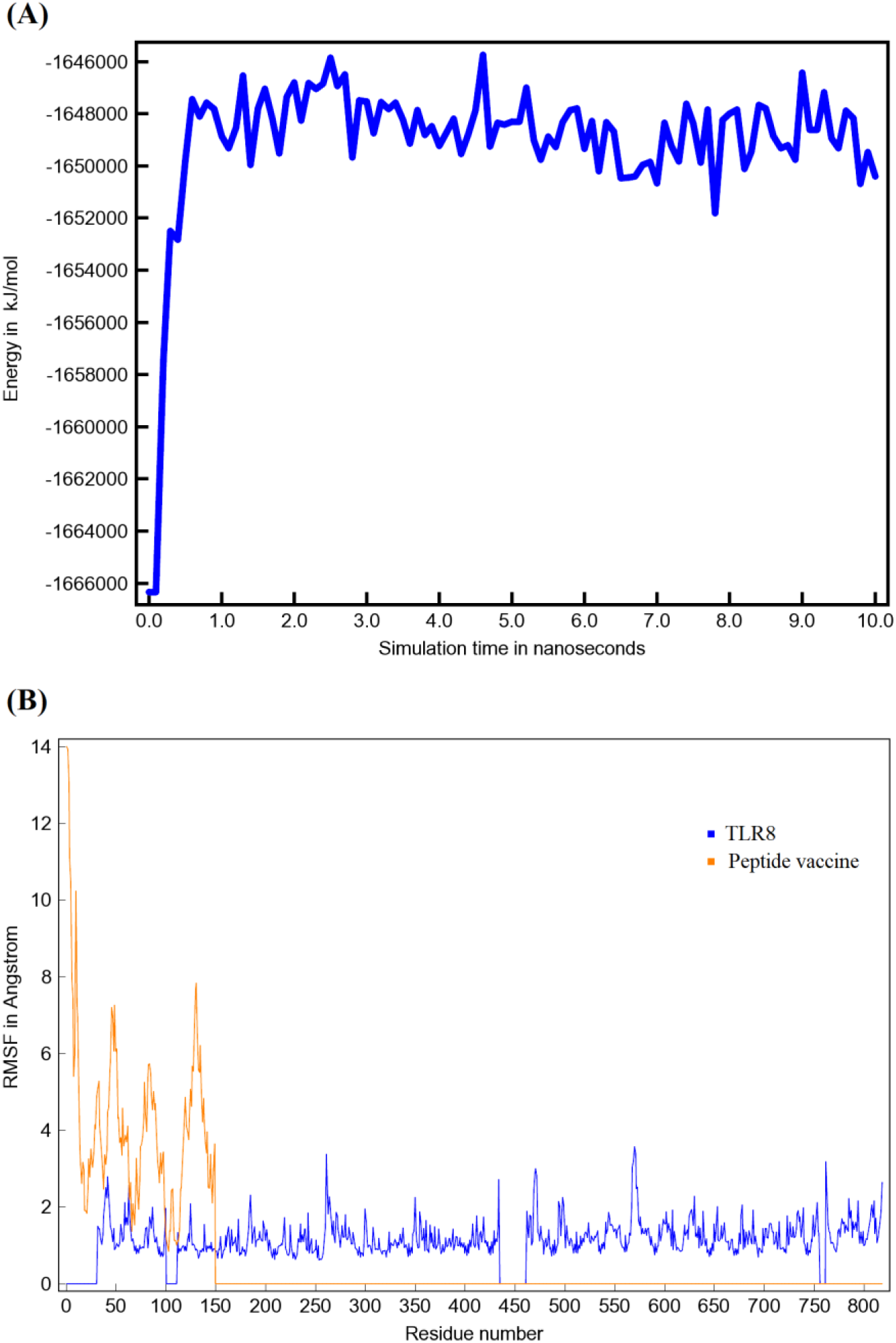
**(A)** Total potential energy of docking complex as a function of simulation period. **(B)** RMSF value calculated for each residue of TLR8 and vaccine construct during simulation. An RMSF value of zero refers to the absence of residue in the molecule under simulation.

Finally, the results of simulation for host immune response against injected vaccine can be seen in **Figure 7**. As predicted by simulation server, we found that three injections of vaccine construct with a time frame of 4 weeks between each two doses are necessary to produce log-lasting and efficient immune response. According to **Figure 7 (A)**, high immunoglobulins titer was generated after the second and third vaccine injections. The titer of various immunoglobulins was kept within a significant level during simulation period even after vaccine level decline. Following vaccine administration, there was a sharp increase in B-cells population including memory B-cells as seen in **Figure 7 (B)**. Based on **Figure 7 (C)**, The B-cells were active during simulation period with an increase in cells duplication and antigen presentation following vaccine injections. The level of helper T-cells, including memory subtype, was significantly high especially after second vaccine injection as presented in **Figure 7 (D)**. As seen in **Figure 7 (E)**, there was an enhancement in macrophages activity and antigen presentation at vaccine injection time-steps. Lastly, there was a remarkable rise in the level of interferon gamma (INF-γ), and interleukin-2 (IL-2) as observed in **Figure 7 (F)**. A sharp elevation in IL-2 level was observed after the second vaccine injection. In summary, these simulation results indicate that the vaccine construct can elicit efficient and long-lasting immune response.

**Figure 7:**
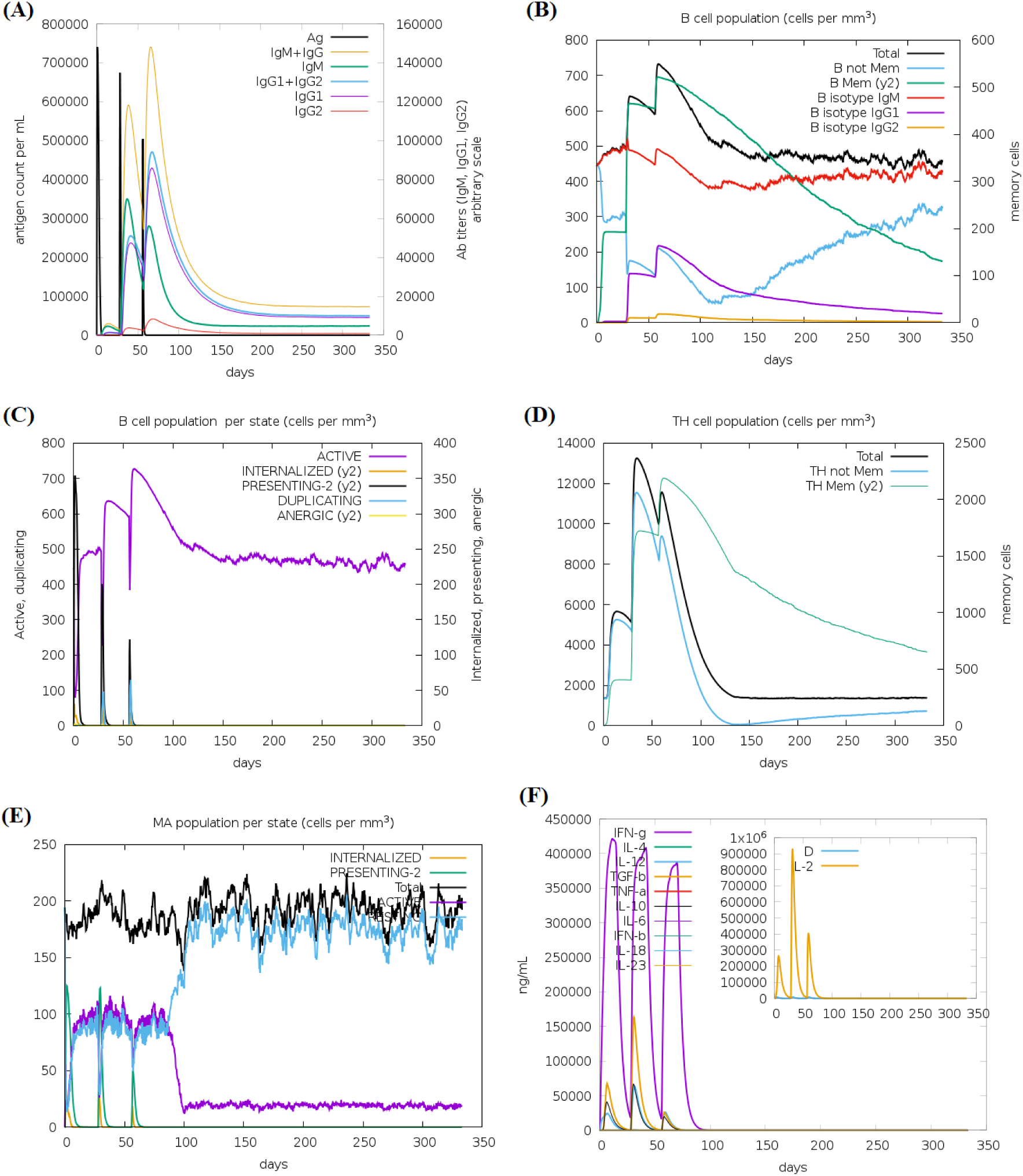
Simulation of level and activity state for various immune system components following vaccine injections. **(A)** Immunoglobulins titer and vaccine level. **(B)** B-cells population level. **(C)** B-cells population per activity state. **(D)** Helper T-cells population level. **(E)** Macrophages population per activity state. **(F)** The levels of interleukins and cytokines expressed as ng/ ml. The D letter in inset plot refers to danger signal.

## Conclusion

Here, we report a peptide vaccine construct with multiple epitopes against SARS-CoV-2 spike protein. The vaccine design seems to be efficient, stable and safe as indicated by prediction, docking and simulation analyses. It may be a potential candidate for further in vitro and in vivo evaluation.

